# Recurrent Interneuron Connectivity does not Support Synchrony in a Biophysical Dentate Gyrus Model

**DOI:** 10.1101/2024.04.16.589667

**Authors:** Daniel Müller-Komorowska, Temma Fujishige, Tomoki Fukai

## Abstract

Synchronous activity of neuronal networks is found in many brain areas and correlates with cognition and behavior. Gamma synchrony is particularly strong in the dentate gyrus, which is thought to process contextual information in the hippocampus. Several network mechanisms for synchrony generation have been proposed and studied computationally. One such mechanism relies solely on recurrent inhibitory interneuron connectivity, but it requires a large enough number of synapses. Here, we incorporate connectivity data of the dentate gyrus into a biophysical computational model to test its ability to generate synchronous activity. We find that recurrent interneuron connectivity is insufficient to induce a synchronous network state. This applies to an interneuron ring network and the broader dentate gyrus circuitry. In the asynchronous state, recurrent interneuron connectivity can have small synchronizing effects but can also desynchronize the network for specific types of synaptic input. Our results show that synchronizing mechanisms relying solely on interneurons are unlikely to be biologically plausible in the dentate gyrus.

## 1 Introduction

Synchronous neuronal activity is important for a range of brain functions. Synchrony can occur at different frequencies, and the gamma range (30 Hz-100 Hz) has been implied by correlation and experiment with cognition and behavior (Kitanishi et al., 2015; Fries, 2009; Wang, 2010; Girardeau et al., 2009). The same implication holds for several brain disorders, such as epilepsy or Parkinson’s disease (Hammond, Bergman, and Brown, 2007; Uhlhaas and Singer, 2006). In the dentate gyrus (DG), gamma oscillations are particularly strong (Buzsáki, Vanderwolf, et al., 1983). Several network mechanisms that could generate these oscillations have been proposed, the interneuron gamma generator (ING) being one of them.

Experiments show that inhibitory interneurons (INs) have a central role in the generation of gamma oscillations (Bartos, Vida, and Jonas, 2007; Allen and Monyer, 2015; Buzsáki and Wang, 2012; Wulff et al., 2009; Antonoudiou et al., 2020). Two major circuit models could allow INs to generate synchrony: the ING model and the pyramidal-interneuron gamma (PING) model (Tiesinga and Sejnowski, 2009). In the ING model, INs synchronize themselves during asynchronous input solely through recurrent inhibition (Wang and Buzsáki, 1996; Bartos, Vida, Frotscher, et al., 2002). In the PING model, on the other hand, the network becomes synchronized through the reciprocal connections of principal cells and INs. The ING and the PING models both work well in-silico and have shown merit in explaining experimental results in some brain areas. However, the ING mechanism requires sufficiently many recurrent inhibitory connections, where the necessary amount depends on intrinsic and synaptic properties (Wang and Buzsáki, 1996; Golomb and Hansel, 2000). Therefore, determining whether the ING mechanism is biologically plausible in a given area requires a careful analysis of the interplay between connectivity and intrinsic properties.

The DG is a hippocampal brain region that receives input primarily from the entorhinal cortex (EC) and sends its output to CA3 (Amaral, Scharfman, and Lavenex, 2007). It is essential for pattern separation (Yassa and Stark, 2011) and has the largest gamma power among the hippocampal regions (Buzsáki, Vanderwolf, et al., 1983). The DG has been proposed to act as a gamma generator. However, unlike the CA3 generator, it is not independent of EC input (Csicsvari et al., 2003). Since the EC is already gamma frequency modulated, it is unclear to what extent the DG generates its own gamma rhythm or whether it merely relays EC gamma. At least at theta frequencies, the EC strongly drives the DG (Mizuseki et al., 2009). The distinction between internally and externally generated oscillations is crucial to restore healthy brain rhythms.

IN types and circuitry of the DG have been extensively studied, and parvalbumin-positive (PV^+^) INs in particular, have revealed extensive recurrent connections (both electrical and chemical) with other PV^+^ INs (Espinoza et al., 2018). This connectivity, together with their intrinsic properties and their fast signaling (Bartos, Vida, Frotscher, et al., 2002), in theory, makes them an excellent substrate for an ING mechanism. However, optogenetic experiments have not been performed on DG PV^+^ INs while measuring synchronous activity and such experiments would not distinguish between ING and PING mechanisms. In practice, it is therefore unclear whether PV^+^ IN connectivity supports synchrony generation. We have thus incorporated the PV^+^ IN connectivity as measured by Espinoza et al., 2018 into a biophysical DG model to study its ability to generate synchronous activity.

Here, we use a biophysical computational network model of the DG (Braganza et al., 2020; Müller-Komorowska et al., 2023) with connectivity data of the PV^+^ INs in the DG of mice (Espinoza et al., 2018) and find that an ING mechanism is unlikely to support synchronous activity. Our results predict that experimental inhibition of recurrent PV^+^ IN connectivity would have little to no effects on synchronous activity. This highlights that even circuit motifs that are qualitatively good candidates to support a specific network dynamic might be insufficient due to quantitative features.

## 2 Methods

### 2.1 PV^+^ Interneuron Ring Model

We used a biophysical parvalbumin-positive (PV^+^) interneuron (IN) model as well as the inhibitory PV^+^-PV^+^ synapse parameters that are described in Braganza et al., 2020. However, our previous model did not have continuous distance-dependent connection probabilities for chemical synapses and did not model gap junctions (GJ).

To model connection probabilities we fit sigmoid functions to data in Espinoza et al., 2018 for chemical synapses as well as GJs (**Supp. Fig. 1**). To model the distance between neurons, we distributed them equidistantly on a ring with a circumference of 4 mm and calculated intersomatic distance on the ring. For our main simulations, we chose to model 120 PV^+^ INs, which is roughly the number of PV^+^ INs in a 300 µm coronal slice of rat hippocampus. The entire unilateral rat dentate gyrus (DG) contains about 4000 PV^+^ INs (Huusko et al., 2015) and it is about 1 cm in length from septal to temporal tip. We chose 120 PV^+^ INs for most simulations because our DG model is the model of a dorsal DG slice and upscaling the model to accommodate 4000 PV^+^ INs would have made it extremely slow to simulate. Self-connectivity was excluded. Random realizations of this connectivity are shown in **Figure 1D**. To model GJs we used the NEURON mechanism from Fernanda Saraga, Ng, and Skinner, 2006. The resistance parameter was set manually to result in a plausible coupling coefficient of about 0.01, measured from soma to soma (**Figure 1B**). The GJ mechanisms were placed in the middle of a randomly chosen proximal dendritic segment, about 32.5 µm from the middle of the soma. To bring the network into an asynchronous state, each neuron was injected at the soma with a constant current randomly chosen from a normal distribution with *I*_*µ*_ = 300 pA and *I*_*σ*_ = 50 pA. *I*_*µ*_ was gradually increased in **Figure 3** to increase the overall network activity. For simulations of PV^+^ INs with *I*_*h*_ we used a NEURON mechanisms first described in Saraga et al., 2003. We added this mechanism to the soma of our PV^+^ IN model and adjusted the channel density parameter to result in a sag amplitude similar to that of CA3 PV^+^ INs (Papp et al., 2013). **Supp. Fig. 3A** shows the properties of the PV^+^ IN with *I*_*h*_.

**Figure 1:**
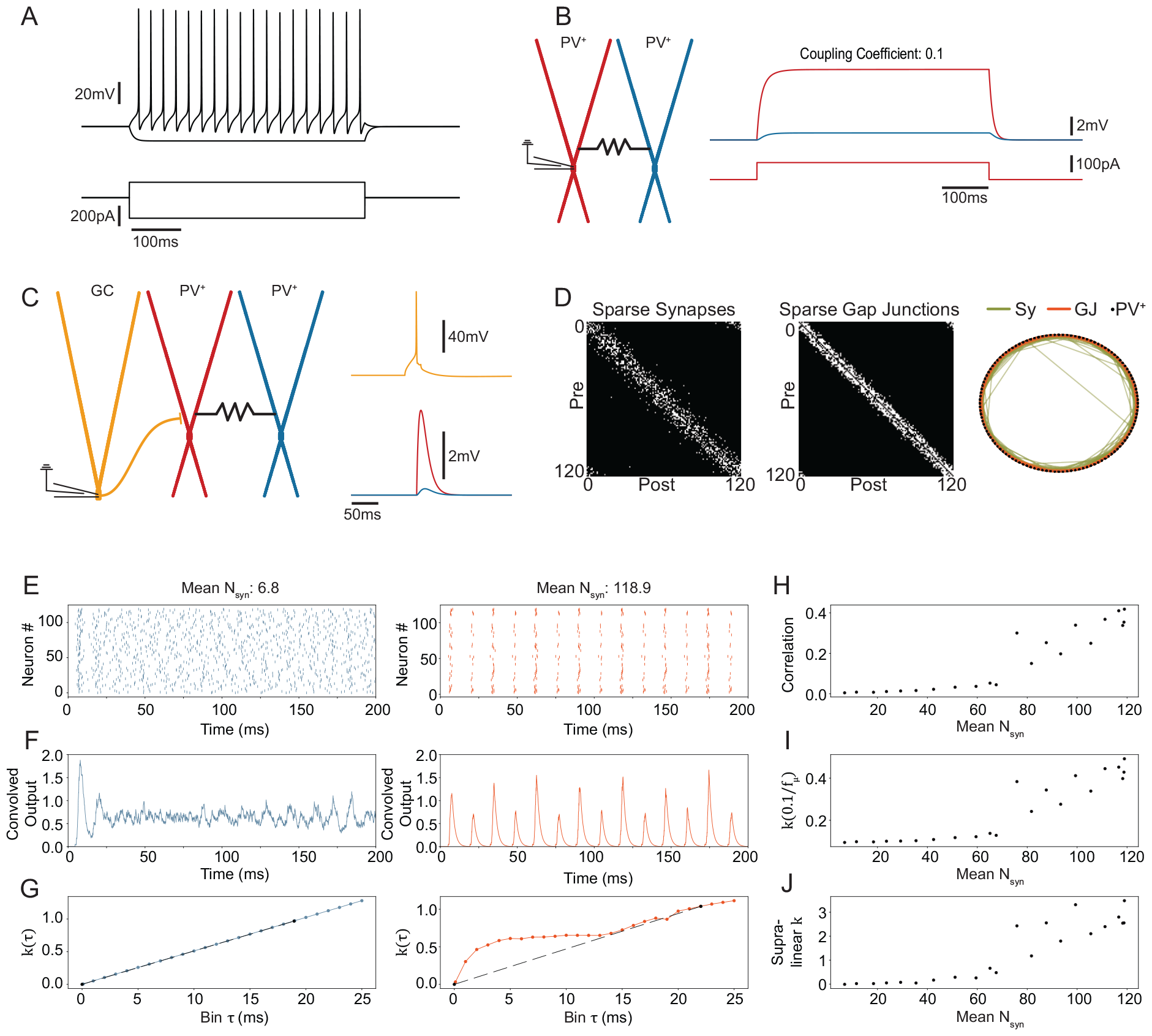
Dense recurrent connectivity creates a synchronous network state but biologically plausible connectivity is insufficient. **A** Intrinsic properties of the biophysical PV^+^ IN model. Note that it does not have *I*_*h*_, as seen in the hyperpolarizing current step. **B** Gap junction coupling implementation. The current injected in the red neuron leads to a somatic voltage drop in the blue neuron. The coupling coefficient is measured at the steady state soma-to-soma. Gap junction resistance was hand-tuned to lead to the coupling coefficient. The gap junction was placed in the middle of the proximal dendritic segment (32.5 µm from the center of the soma). **C** Granule cell EPSP is measurable in a coupled PV^+^ IN. Gap junction resistance and placement as in B. **D** Random realization from the sigmoid functions fit to the measured connection probabilities from Espinoza et al., 2018 and the distances of the ring network (see methods). **E** Spike raster plots of 120 PV^+^ INs with biologically plausible connectivity on the left in blue (D shows a biologically plausible connectivity example) and dense (nearly full) chemical connectivity on the right in orange. *N*_*syn*_ is the number of chemical synapses received by the average neuron in this example run. **F** The convolved output is calculated from the spike rasters above by convolving with a synaptic kernel. Shows the synchronous state for the full connectivity on the right. **G** The coherence measure *k*(*τ*) for different values of *τ* (see methods) for the biologically plausible on the left (blue) and the dense connectivity on the right (orange). The dashed black line shows the line between (0, 0) and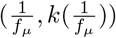, where *f*_*µ*_ is the average frequency of all neurons. **H-J** Synchrony measures at different synaptic densities. All three show a non-linear increase of around 60-70 synapses. **H** is the average pairwise correlation coefficient, **I** is the coherence measure *k* calculated as the inverse of the average network frequency and **J** is the estimated area between the line and the actual function of *k*(*τ*) (see G). PV, parvalbumin; GC, granule cell; Sy, synapse; GJ, gap junction.

### 2.2 Large Scale PV^+^ IN Plane Model

To check whether the effects of IN connectivity on synchrony generation differ depending on the size of the network, we simulated 4000 PV^+^ INs, since that represents roughly their total number in an entire unilateral DG. We also change from a ring network topology to a plane. The 2D plane was 1 cm in length and 0.4 cm in width. Cells were distributed uniformly on it. Distance-dependent connectivity was maintained as above through sigmoid functions. The results are shown in**Supp. Fig. 2**.

**Figure 2:**
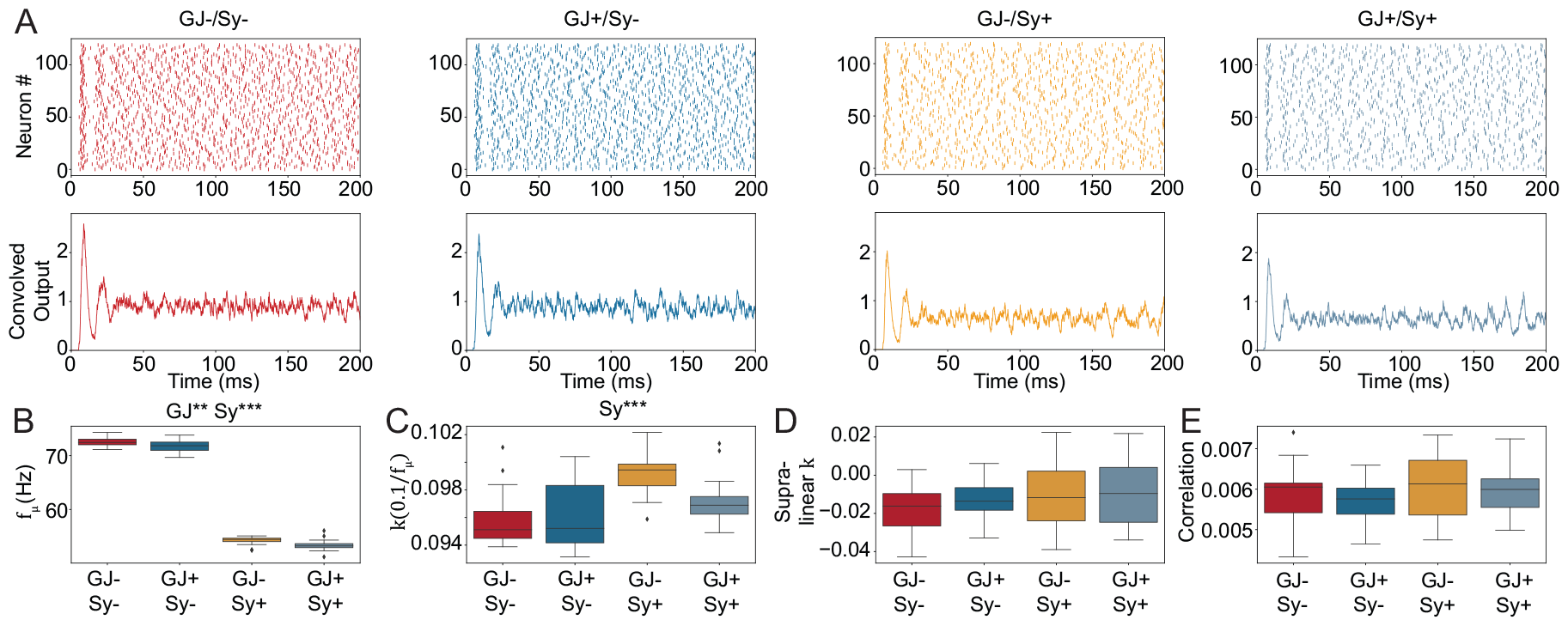
Adding PV^+^ IN connectivity to the ring network does not consistently increase network synchrony. **A** Spike raster plots (top) and convolved output (bottom) for three different conditions of the PV^+^ ring network. GJ-/Sy-has neither gap junctions nor chemical synapses. GJ+/Sy-has only gap junctions, GJ-/Sy+ has only chemical synapses and GJ+/Sy+ has gap junctions and chemical synapses. GJ+/Sy+ is a different random realization of the connectivity shown in **Figure 1D**. The convolved output does not show a synchronous state. **B** Average frequency including all neurons of the network. Both gap junctions and chemical synapses significantly decrease the average activity of the ring network with a much larger effect of chemical synapses. Two-way ANOVA: Interaction: *F* = 0.0133, *p* = 0.9084, Main effects: GJ, *F* = 8.7578, *p <* 0.01; Sy, *F* = 5206.6925, *p <* 0.001. **C-E** The three synchrony measures. Two-way ANOVA only showed significant effects for 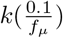 (C): *F* = 3.7789, *p* = 0.0569, Main effects: GJ, *F* = 2.2704, *p* = 0.1375; Sy, *F* = 15.8746, *p <* 0.001. For each condition ten samples were simulated with different random seeds. ^*∗*^*p <* 0.05, ^*∗∗*^*p <* 0.01, ^*∗∗∗*^*p <* 0.001, absence of asterisk indicates *p >*= 0.05, statistically insignificant.

**Figure 3:**
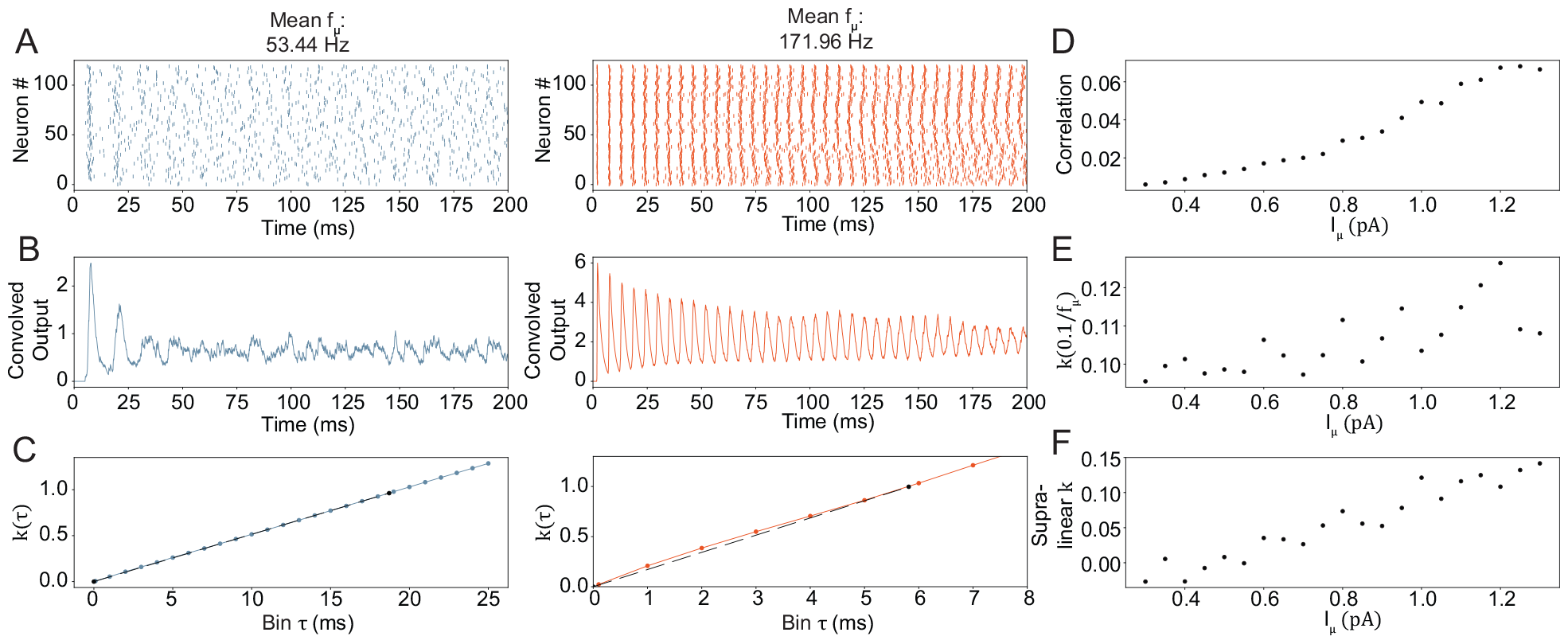
Increasing the input drive increases network synchrony but does not reach the magnitude found with dense network connectivity. **A** Spike raster plots for the PV^+^ ring network with biologically plausible connectivity for the standard model on the left (blue, input strength as in **Figure 1,2**), and a model with strong input drive on the right. The mean frequency *f*_*µ*_ is calculated from all neurons in the network. **B** The convolved output of the above spiking activity. **C** The coherence measure *k*(*τ*) for different values of *τ* (see methods) for the baseline activity condition on the left (blue) and the high activity on the right (orange). The dashed black line shows the line between (0, 0) and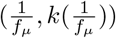, where *f*_*µ*_ is the average frequency of all neurons. **D-F** Synchrony measures at different synaptic densities. All three show increasing synchrony with increasing input current. However, note that none of them reaches the synchronous regime shown in **Figure 1H-I**. H is the average pairwise correlation coefficient, I is the coherence measure *k*(*τ*) calculated at the inverse of the average network frequency and J is the estimated area between the line and the actual function of *k* (see G). The connectivity in these simulations is biologically plausible. The independent parameter *I*_*µ*_ is the mean of the input strength distribution.

### 2.3 The PV^+^ IN Ring Network in the Larger DG Model

We integrated the PV^+^ IN ring model with distance-dependent chemical and GJ connectivity into our larger DG model (Braganza et al., 2020; Müller-Komorowska et al., 2023). To accommodate 120 PV^+^ INs we scaled up the model by a factor of five to 10000 granule cells (GCs), 300 mossy cells (MCs), 120 PV^+^ INs, and 120 hilar-perforant path associated cells (HCs). We removed the input current that generated asynchrony in the above versions of the ring model and instead added Poisson input processes that model input from 120 entorhinal cortex cells. For the homogeneous Poisson input (**Figure 4**) the average input rate was 15 Hz. For the gamma oscillation conditions (**Figure 5**) an inhomogeneous Poisson process was used with a sinusoidal waveform at 30 Hz (slow gamma) or 80 Hz (fast gamma). For **Figure 6** we additionally recorded the somatic membrane voltage of all granule cells.

**Figure 4:**
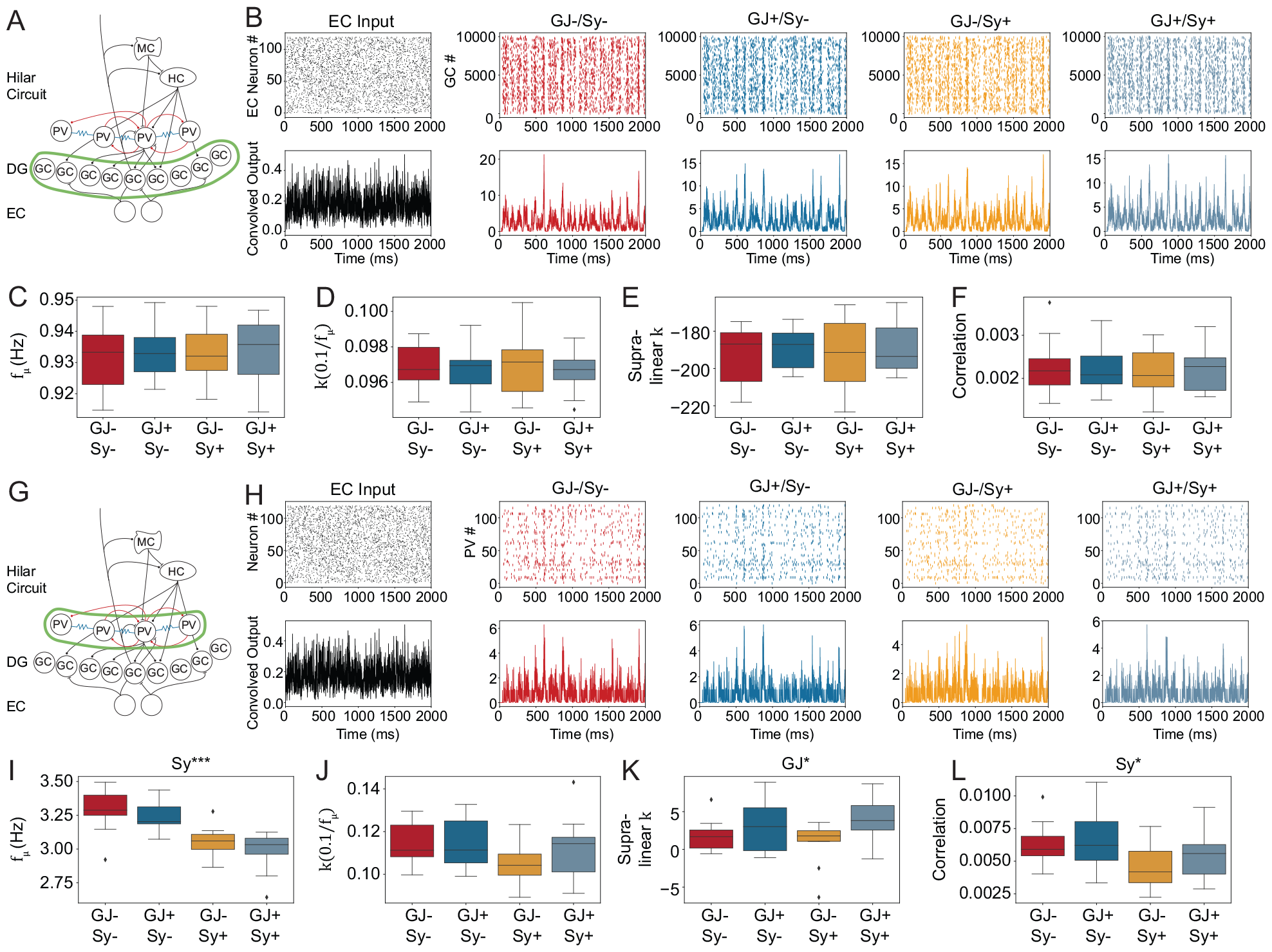
The PV^+^ IN ring network connectivity has no effect on GC synchrony and inconsistent effects on the PV^+^ ring network itself in the biophysical DG model. **A**,**G** Schematic of the DG model. Green highlights the cells that are analyzed. B-F, GCs and H-L PV^+^ cells. Only the connectivity of the ring network was changed in the different conditions. **B** Spike raster plots (top) and corresponding convolved output of EC neurons (black) and GCs. Note that the time constant of the kernel to calculate the convolved output was kept constant to the fast 1.8 ms of the PV^+^ neuron synapse for all cell types. **C** Average frequency of the network calculated from all neurons. Two-way ANOVA showed no significant effects. **D-F** The synchrony measures. Two-way ANOVA showed no significant effects. Note that in E values are large and negative. This is likely a failure of the supralinear k measure that occurs when the average frequency is slow. **H** Same as B but with spike raster plots and convolved output for the PV^+^ INs instead of the GCs. **I** Average frequency of the network calculated from all neurons. Two-way ANOVA: Interaction: *F* = 0.0881, *p* = 0.7683, Main effects: GJ, *F* = 1.8815, *p* = 0.1787; Sy, *F* = 30.9370, *p <* 0.001. **J-L** The synchrony measures as in D-F but for the PV^+^ neurons. Two-way ANOVA showed no significant results in J. Two-way ANOVA for Supralinear *K*: Interaction: *F* = 0.9901, *p* = 0.3264, Main effects: GJ, *F* = 5.7611, *p <* 0.05; Sy, *F* = 0.0054, *p* = 0.9421. Two-way ANOVA for Correlation: Interaction: *F* = 0.2310, *p* = 0.6337, Main effects: GJ, *F* = 0.7404, *p* = 0.3952; Sy, *F* = 5.5301, *p <* 0.05.

**Figure 5:**
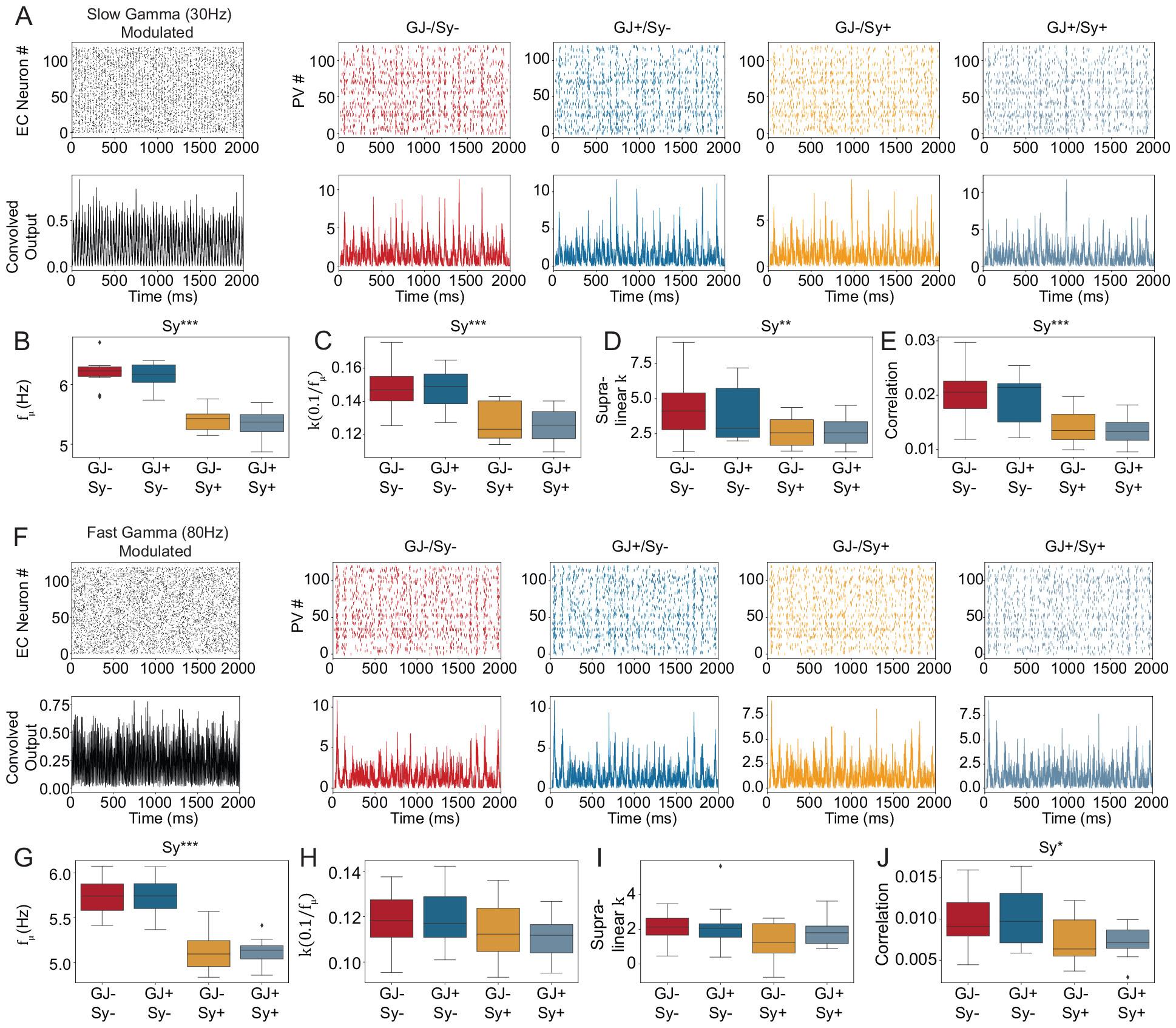
Recurrent Connectivity desynchronizes the PV^+^ ring network when the input is 30 Hz frequency modulated but not when the input is 80 Hz modulated. This figure’s shows results from the biophysical DG model but results for the other cell types are shown in a supplemental figure. **A** Spike raster plots (top) and corresponding convolved output of EC neurons (black) and GCs in the slow gamma (30 Hz) modulated condition. Note that the time constant of the kernel to calculate the convolved output was kept constant to the fast 1.8 ms of the PV^+^ neuron synapse for all cell types. **B** Average frequency of the network calculated from all neurons. Two-way ANOVA: Interaction: *F* = 0.0215, *p* = 0.8842, Main effects: GJ, *F* = 0.0256, *p* = 0.8737; Sy, *F* = 80.9939, *p <* 0.001. **C-E** The synchrony measures for the PV^+^ neurons. Two-way ANOVA for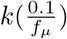 : Interaction: *F* = 0.0392, *p* = 0.8440, Main effects: GJ, *F* = 0.0586, *p* = 0.8101; Sy, *F* = 28.619, *p <* 0.001. Two-way ANOVA for Supralinear *K*: Interaction: *F* = 0.3101, *p* = 0.5811, Main effects: GJ, *F* = 0.2058, *p* = 0.6528; Sy, *F* = 7.9659, *p <* 0.01. Two-way ANOVA for Correlation: Interaction: *F* = 0.000659, *p* = 0.979662, Main effects: GJ, *F* = 0.406382, *p* = 0.527845; Sy, *F* 20.858620, *p <* 0.001. **F** Spike raster plots (top) and corresponding convolved output of EC neurons (black) and GCs as in A but for the fast gamma (80 Hz) condition. **G** Average frequency of the network calculated from all neurons. Two-way ANOVA: Interaction: *F* = 0.0215, *p* = 0.8842, Main effects: GJ, *F* = 0.0256, *p* = 0.8737; Sy, *F* = 80.9939, *p <* 0.001. **H-J** The synchrony measures for the PV^+^ neurons in the fast gamma condition. Two-way ANOVA for 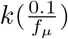 and supalinear *k* was insignificant. Two-way ANOVA for Correlation: Interaction: *F* = 0.1609, *p* = 0.6907, Main effects: GJ, *F* = 0.00003, *p* = 0.9952; Sy, *F* = 7.0808, *p <* 0.05.

**Figure 6:**
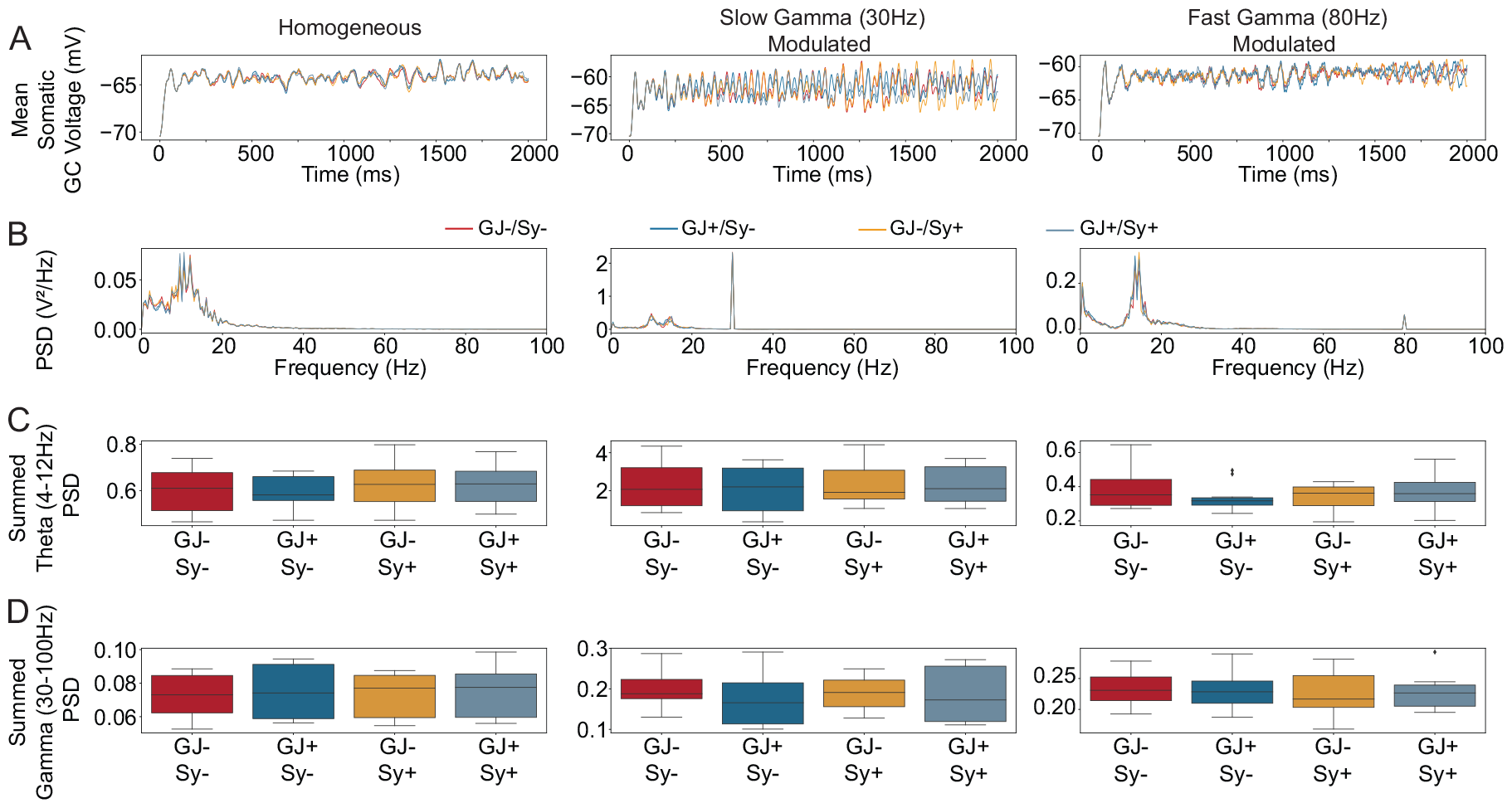
Inhibitory recurrence does not change theta or gamma power measured through the GC membrane voltage. **A** Single run examples of the average membrane potential across all granule cells. The color labeling for the conditions is between A and B and applies to both panels A & B. **B** The power spectral density of the average membrane potential averaged across ten different simulation runs per condition. **C** Boxplots of the summed PSD in the theta range. Two-way ANOVA showed no significant main effects or interactions for any of the conditions. **D** Boxplots of the summed PSD in the gamma range. Two-way ANOVA showed no significant main effects or interactions for any of the conditions.

### 2.4 Network Synchrony Measures

We mainly use the *k*(*τ*) measure (Wang and Buzsaki, 1996) to quantify the overall network synchrony. *k*(*τ*) is calculated as the cross correlation between two binned spike trains. The spike trains can be binned at different bin sizes *τ*. Given the total length of the simulation *T* and bin size *τ* we have 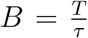 bins. An incomplete bin at the end is discarded. The coherence measurement between the spike trains of two neurons *i, j* and their binned spike trains *b*_*i*_ and *b*_*j*_ is:

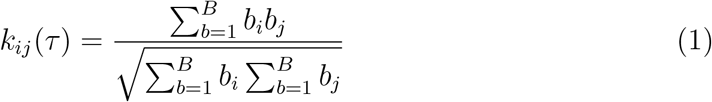

To calculate *k*(*τ*) we take the mean of *k*_*ij*_ for all unique pairs of neurons where neither of the neurons is inactive during the trial and *i* ≠ *j*:

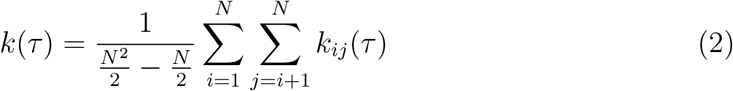

Between 0 *< τ < f*_*µ*_, *k*(*τ*) increases linearly with *τ* if the network is asynchronous. *f*_*µ*_ is the average frequency of the network’s cells. In a synchronous network *k*(*τ*) will be supralinear in some places. Since the slope of the line also depends on *f*_*µ*_, the coherence measure needs to be frequency corrected. **We therefore use the measure** 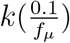 **to quantify synchrony**. This measure is used in Bartos, Vida, Frotscher, et al., 2002. However, the choice of 0.1 in the numerator is arbitrary. We therefore also measured synchrony by estimating the area between the line that goes through (0, 0) and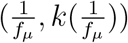, and the curve *k*(*τ*) from 0 up to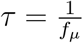. The estimation is done by calculating *k*(*τ*) for 30 equidistant values of *τ* between 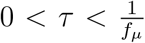. **We call this measure supralinear k** because it represents the extent to which *k*(*τ*) is above the line that would be expected from an asynchronous network. We also calculated the correlation coefficient of the convolved output from each neuronal pair where both neurons fire at least one action potential. The convolved output was calculated by convolving a neuron’s spike train with an exponential decay function with a time constant of 1.8ms. This is the time constant of synaptic decay of a PV^+^ IN synapse in the model. **The correlation synchrony measure is the average correlation coefficient of the convolved output between all valid neuron pairs**. The convolved outputs that are plotted were calculated by convolving the summed spike trains of all neurons in a population with the same exponential decay function.

Finally, to quantify the power at different frequencies (**Figure 6**) we calculated the mean membrane voltage of all GCs. We then estimated the power spectral density (PSD). For statistical testing the PSD at theta (4 Hz-12 Hz) and gamma (30 Hz-100 Hz) was summed up to get a single number for power in each frequency band.

### 2.5 Software and Simulation

Simulations were done with NEURON 8.0.0 (Carnevale and M. L. Hines, 2006) in Python 3.9.12 (M. Hines, Davison, and Muller, 2009). Plotting was done with matplotlib 3.5.3 (Hunter, 2007) and seaborn 0.11.2 (Waskom, 2021). Tabular data was organized with pandas 1.4.3 (Reback et al., 2022). The statistical test to assess the effect of chemical synapses and GJs was the two-way ANOVA as implemented in the statsmodels 0.13.2 Python package (Seabold and Perktold, 2010). To calculate the PSD we used the periodogram function from the scipy signal toolbox (Virtanen et al., 2020). *p* values were considered significant if *p <* 0.05. *p* and *F* values were rounded to the fourth significant decimal. For each condition, ten different networks were sampled using different random seeds. The Δ*t* of all simulations was 0.0001 s. The duration of all simulations was 2 s and the entire duration was used in the analysis.

## 3 Results

### 3.1 Synchrony Emerges from Dense Recurrent Inhibition

To investigate the ability of biologically plausible connectivity to generate synchrony we started with a ring network consisting solely of parvalbumin-positive (PV^+^) interneurons (INs). We used the biophysical model of a dentate gyrus (DG) PV^+^ IN that we used previously as part of a DG network model (Braganza et al., 2020; Müller-Komorowska et al., 2023). This model exhibits continuous firing in response to somatic step current injections and does not contain *I*_*h*_ channels as seen from the lack of slow depolarization in the hyperpolarizing current step (**Figure 1A**). To model the biologically plausible connectivity, we used paired patch clamp data from Espinoza et al., 2018. For gap junctions, they measured no distance dependence of the coupling coefficient. Therefore, we gave all gap junctions the same resistance to result in a coupling coefficient of about 0.1 measured from soma to soma (**Figure 1B**). For lack of DG specific anatomical data, gap junctions were placed in the middle of the proximal dendritic segment, about 32.5 µm from the center of the somatic segment. We also ensured that a granule cell EPSP (as modeled in Braganza et al., 2020) is measurable at the soma of a coupled PV^+^ IN (**Figure 1C**), a phenomenon measured by Espinoza et al., 2018.

To model the connection probability we hand-fit sigmoid functions to resemble the somatic distance-connection probability functions reported in Espinoza et al., 2018 for both gap junctions and chemical synapses (**Supp. Fig. 1**). To model the overall number of PV^+^ INs and their somatic distance we assumed that a unilateral DG contains about 4000 PV^+^ INs (Huusko et al., 2015). Furthermore, to represent a 300 µm coronal slice we simulate 120 PV^+^ INs assuming that the DG is about 1 cm from temporal to medial tip in the rat. To simplify anatomy we assumed a ring where neurons are distributed equidistantly, resulting in 33.3 µm between neurons. **Figure 1D** shows an example of a random realization of the sigmoidal connection probabilities and the neuronal distance. We took chemical synapses’ strength and decay time constant from our previous model implementation.

With this ring model in place, we moved to investigate synchrony in the spiking output of the network. To induce asynchronous activity each neuron receives a current injection randomly drawn from a normal distribution. We found that for the biologically plausible connectivity, where the average PV^+^ IN receives 6.8 chemical synapses, the network remains asynchronous. However, when the sigmoid function is adjusted for nearly full chemical connectivity (118.9 synaptic inputs per neuron), the network is synchronous (**Figure 1E,F**). To quantify synchrony, we calculate the measure *k*(*τ*) on the binned spike trains (**Figure 1G**, see methods). Furthermore, we also calculate the average correlation coefficient of each neuronal pair in the network (**Figure 1H**). All synchrony measures show that there is a discontinuity in network state at about 60-70 synapses per neuron. At sparser connectivity, synchrony increases only slightly with increasing synapse number. At about 60-70 synapses, synchrony changes slope and then increases linearly. This suggests that at dense connectivity with about 60-70 synapses, the network enters a synchronous state. At sparse biological connectivity (6.8 synapses) on the other hand network activity is asynchronous. Next we selectively turned the biologically plausible gap junction and chemical connectivity on to investigate their effects on synchrony separately.

### 3.2 Biologically Plausible Recurrent Connectivity is Insufficient for Synchronous State

Neither gap junctions, nor chemical synapses nor their interaction, are sufficient to induce the synchronous state (**Figure 2A**). However, both have significant effects on the average firing rate of the network (**Figure 2B**). Statistical analysis of the three synchrony measures (**Figure 2C-E**) showed a significant increase only of 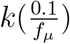 from synaptic connectivity but not from gap junctions or their interaction. However, the magnitude of the effect is small and the synchrony measure is still nowhere near the synchronous state (compare **Figure 1I** to **Figure 2C**). All other tested effects were non-significant.

So far we simulated networks of 120 PV^+^ INs that represent a small part of the entire DG PV^+^ population. To ensure that we do not miss the effects of network size, we simulated networks with 4000 PV^+^ INs with somatic positions randomly distributed on a 2D plane. This also changes the originally equidistant anatomical arrangement of the cells. In this large network, we again found no synchronous state, despite clear effects of both gap junctions and chemical synapses on the average rate (**Supp. Fig. 2**).

Although no strong I_*h*_ dependent sag potential has been measured in DG PV^+^ INs (Lübke, Frotscher, and Spruston, 1998), could *I*_*h*_ be necessary in other areas to enable the synchronizing effects of connectivity? To test this we added I_*h*_ to the soma of the PV^+^ model. The result was the same: an asynchronous network. The significant effect of gap junctions on the average firing rate, however, was abolished by *I*_*h*_ (**Supp. Fig. 3**). Within the asynchronous regime, synaptic connections caused small but significant increases in supralinear *k* and the correlation measure. The input drive of the neurons was the same in the models thus far. Because overall activity can influence synchrony, we next simulated different input strengths.

To increase the overall network activity, we increased the mean of the distribution (*I*_*µ*_) from which each cell’s input current was drawn. **Figure 3A** shows the baseline case with the same input strength as above on the left (blue, 53 Hz) and an extreme case with an average frequency of 172 Hz on the right (orange). The convolved output shows extremely fast synchrony that decreases over time (**Figure 3B**). Analysis of *k*(*τ*) for different values of *τ* shows only minor deviation from linearity (**Figure 3C**). All three synchrony measures (**Figure 3D-F**) show an increase of synchrony with increasing input strength that does not reach the magnitude of synchrony in the synchronous state as shown in **Figure 1H-J**. To check whether connectivity influences the synchrony measures at high frequency we turned connectivity on and off as above (**Supp. Fig. 4**). This showed that the supralinear *k* and the correlation increase significantly when synaptic connectivity is enabled. This result could have implications for in-vitro conditions, where an excitatory agonist increases the excitatory drive on inhibitory INs. However, a consistent frequency of 172 Hz is not sustainable in-vivo over long time frames. During shorter time windows on the other hand, on the order of a dendritic spike or a gamma burst, this type of high-frequency synchrony could be relevant. During short but strong input the temporal dynamics of the principal cells strongly factor into the activity of INs. We, therefore went on to integrate the ring model into our larger DG model.

In the DG the GCs are the principal cell type and outnumber basket cells by a factor of 1:100 in the suprapyramidal and 1:180 in the infrapyramidal blade (Amaral, Scharfman, and Lavenex, 2007). We therefore first investigated whether connectivity in the PV^+^ ring network affects GC activity (**Figure 4A**). As expected, biologically plausible IN connectivity did not synchronize the GC population (**Figure 4B**). Even the average frequency of GCs is not significantly affected by IN connectivity (**Figure 4C**) and neither are any of the three synchrony measures (**Figure 4D-F**). This is in contrast to the PV^+^ neurons (**Figure 4G**). While they did not reach a synchronous state (**Figure 4H**), their average frequency was significantly decreased by inhibitory synaptic connectivity (**Figure 4I**). The correlation measure of PV^+^ neurons on the other hand was significantly decreased by chemical synapses (**Figure 4L**). However, like the upward trends in the ring network, the effect was small in magnitude and importantly, does not affect GC synchrony. The data for mossy cells and HC cells is shown in **Supp. Fig. 5** but none of the parameters showed statistically significant effects for them. Overall, this data does not give any indication that PV^+^ neurons can generate synchrony given homogeneous Poisson input from entorhinal cortex (EC) neurons. But biologically plausible input is unlikely to be homogeneous. It is rather expected to have some oscillatory structure. We therefore went on to simulate synaptic input that is modulated at slow gamma (30 Hz) or high gamma (80 Hz) frequency.

Because GCs showed no significant effects with the homogeneous Poisson input, we focused this analysis on the PV^+^ INs. For slow gamma (30 Hz) modulated input the PV^+^ INs did not enter a visible synchronous state (**Figure 5A**), although their average frequency was significantly affected by their connectivity (**Figure 5B**). The synchrony measures all showed small but significant decreases in synchrony (**Figure 5C-E**). No significant effects of gap junctions or the interaction were found. For the fast gamma (80 Hz) modulated inputs (**Figure 5F**) the result was overall similar, however 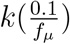 and supralinear *k* were not statistically significant. **Supp. Fig. 6** shows the GCs of the model, which were not synchronized in any of the connectivity conditions. **Figure S 6** shows the other two cell types of the model. Overall we found no indication that recurrent PV^+^ IN connectivity supports the synchronization of the DG model.

So far, we have used generic synchrony measures of the output spike train. During experiments in biological tissue, synchrony is measured through field potential and evaluated at specific frequencies. We therefore finished our analysis by calculating the power spectrum of the average granule cell membrane potentials, because they broadly integrate direct as well as recurrent input currents. We ran simulations for the DG model and the different conditions as shown in the above figures, but recorded the somatic membrane voltage of all 10000 GCs. **Figure 6A** shows the average membrane potential of all GCs of a representative model run for the homogeneous Poisson input, slow gamma (30 Hz) modulated and fast gamma (80 Hz) modulated. Regardless of the input modulation, gap junctions and synapses do not have a clear effect on the oscillatory frequencies. This is also true when looking at the power spectral density (PSD) in **Figure 6B**. This shows the average PSD over 10 model runs in each condition. For statistical testing, we calculated the summed PSD at theta (4 Hz-12 Hz) and gamma (30 Hz-100 Hz). No statistically significant main effects or interactions for any of the conditions were found. This highlights our conclusion that IN recurrence alone does not synchronize the DG network.

## 4 Discussion

We have performed a variety of simulations, all intending to find synchronous neuronal activity and all with the same result: biologically plausible connectivity does not generate synchronous neuronal activity. We started by varying connectivity from sparse and biologically plausible to fully connected. As reported before (Wang and Buzsaki, 1996; Bartos, Vida, Frotscher, et al., 2002), we found that a synchronous state is possible in the ring network if the number of recurrent synapses is large enough. This is consistent with the interneuron gamma generator (ING) model of synchrony. However, more synapses are required than we expect from distance-based connectivity probabilities (Espinoza et al., 2018). Comparing simulations with biologically plausible recurrent connectivity to those without showed small but significant effects for some synchrony measures. Effects were particularly clear when *I*_*h*_ current was added to the neuronal dynamics or the average frequency was high. Nonetheless, synchrony did not reach the same level as we have observed in the synchronous state and was inconsistent across measurements. Furthermore, we observed neither synchrony nor increased theta or gamma power in the larger dentate gyrus (DG) model due to recurrent IN connectivity. On the contrary, PV^+^ INs showed very slight desynchronization for frequency-modulated inputs. Therefore, we conclude that in the DG, recurrent interneuron connectivity is too sparse to cause synchronous activity.

Previous papers most likely found synchrony in ring networks of recurrently connected INs (Wang and Buzsáki, 1996; Bartos, Vida, Frotscher, et al., 2002) because they have assumed more synapses than we have in our model. Theoretical work has well established that in a ring network, synchrony requires a sufficiently large number of inhibitory synapses (Golomb and Hansel, 2000). Below that number, the network is asynchronous. Wang and Buzsaki, 1996 also investigated the influence of synapse number and found that at least 60 synaptic inputs per neuron are required for synchrony. Our estimate of 70 synapses is not far off, although we did not attempt to replicate the exact neuron model, which is known to be relevant to synchronization (Nomura, Fukai, and Aoyagi, 2003). Wang and Buzsaki, 1996 consider 60 a sparse and plausible number based on an estimation of synaptic contacts made by a biocytin filled PV^+^ basket cell in CA1 Sik et al., 1995 and anatomical data regarding PV^+^ neuron density in CA1 (Aika et al., 1994). Bartos, Vida, Frotscher, et al., 2002 extrapolate these CA1 estimates to the DG and thereby likely overstate the number of recurrent DG synapses. The primary difference between ours and previous studies is that we use DG connectivity estimates (Espinoza et al., 2018) instead of CA1 estimates.

The circuit mechanisms of synchrony can also be studied experimentally. Unfortunately, experiments that inhibit PV^+^ INs are inadequate for conclusions about recurrent inhibition as they perturb the influence of all inputs, not only inhibitory recurrence. A study by Wulff et al., 2009 has come close to directly testing the effect of inhibitory recurrence by deleting the GABA_*A*_ *γ*_2_ subunit in PV^+^ INs. Measuring in CA1, they found an effect on gamma oscillations. We predict the same result for the DG. However, Wulff et al., 2009 find effects on theta power, which our model does not predict for the DG. Moreover, the genetic deletion approach has two important limitations: it affects all PV^+^ neurons of the brain and affects mice throughout their development. Therefore, compensatory effects could obscure the role of recurrent inhibition in gamma generation. An opto-/chemogenetic approach targeting recurrent synapses specifically could overcome these limitations, but it is currently infeasible. Alternatively, an inducible genetic deletion could achieve the goal. However, the genetic deletion results from Wulff et al., 2009 support our prediction that recurrent inhibition has no major role in gamma generation.

Besides recurrent inhibition, feedback between pyramidal cells and interneurons can also generate synchrony. This is called the pyramidal-interneuron gamma generator (PING Tiesinga and Sejnowski, 2009) model. Since hippocampal networks feature IN to IN and PC to IN connectivity, both mechanisms are assumed to cooperate in gamma generation (Buzsáki and Wang, 2012). When we turn recurrent connections between INs off, we find no effect on theta or gamma power, suggesting that the two mechanisms do not meaningfully interact. However, while our model has granule cell (GC) to IN feedback connectivity, it does not show gamma power. Therefore, our model does not act as a gamma generator and may not be a good representative of a PING model. For example, our GC model does not exhibit bursting, a known GC dynamic in-vitro and in-vivo (Pan and Stringer, 1996; Pernia-Andrade and Jonas, 2014; Neubrandt et al., 2018). GC bursting could be a necessary component of a DG PING model. In its current implementation, however, recurrent inhibition affects neither theta nor gamma power.

Our major prediction is that selectively disabling recurrent IN connectivity in the DG does not affect synchronous activity. This prediction could become testable as neuroscience develops tools to manipulate smaller and smaller subcellular circuit components. Even circuit components that seem like qualitatively plausible mechanisms can yield negative results based on quantity.

## 5 Acknowledgments

We thank Joanna Komorowska-Müller for her helpful comments on the manuscript. This work was partly supported by JSPS KAKENHI no. JP23H05476.

## 6 Author Contributions

DMK has designed the study, written the code to run the simulations, analyzed the data, prepared the figures and written the initial draft of the manuscript. TFujishige has written code to incorporate the gap junctions into the DG model. TFukai has helped with the study design. All authors have read and contributed to the manuscript.

## 7 Data and Code Availability

The code will be available as a release in the pydentate repository. Data will be made available on Zenodo.

---

**Supp. Fig. 1:**
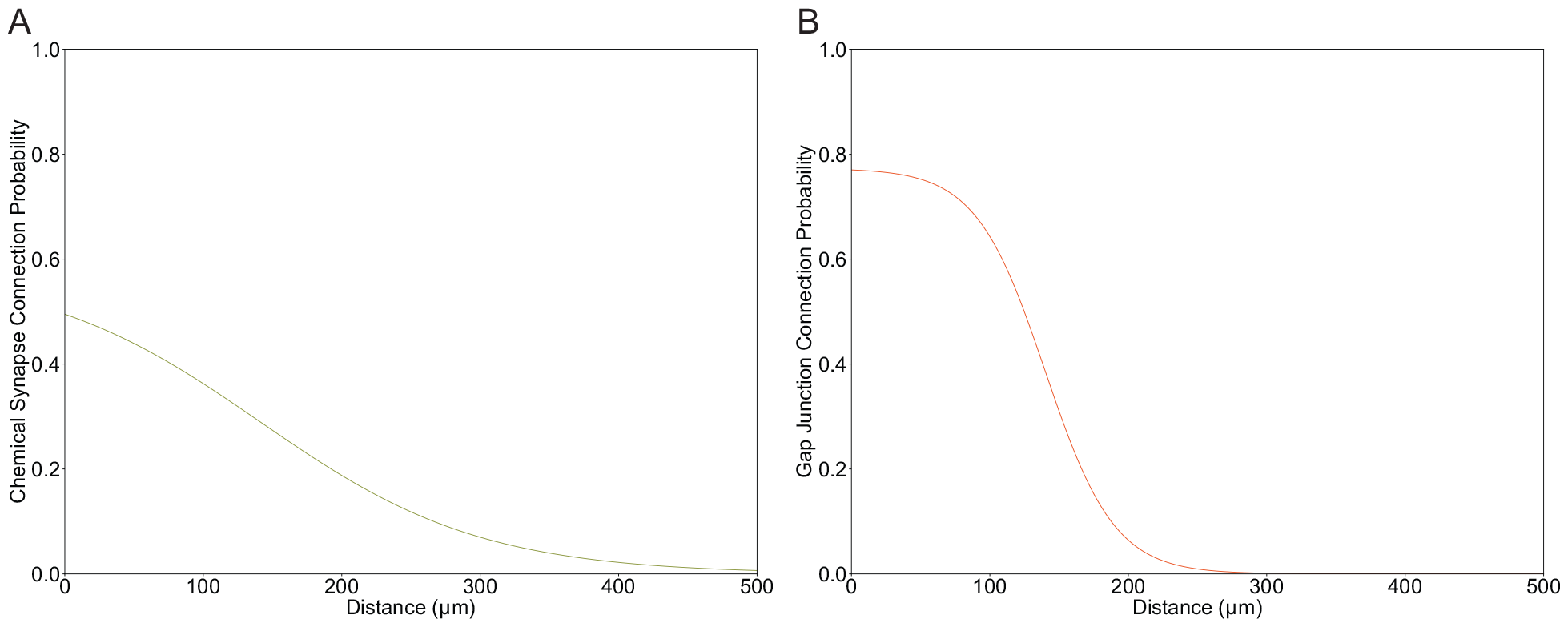
The biologically plausible sigmoid connection probability functions for chemical synapses (A) and gap junctions (B). These were designed to fit the data in Espinoza et al., 2018.

**Supp. Fig. 2:**
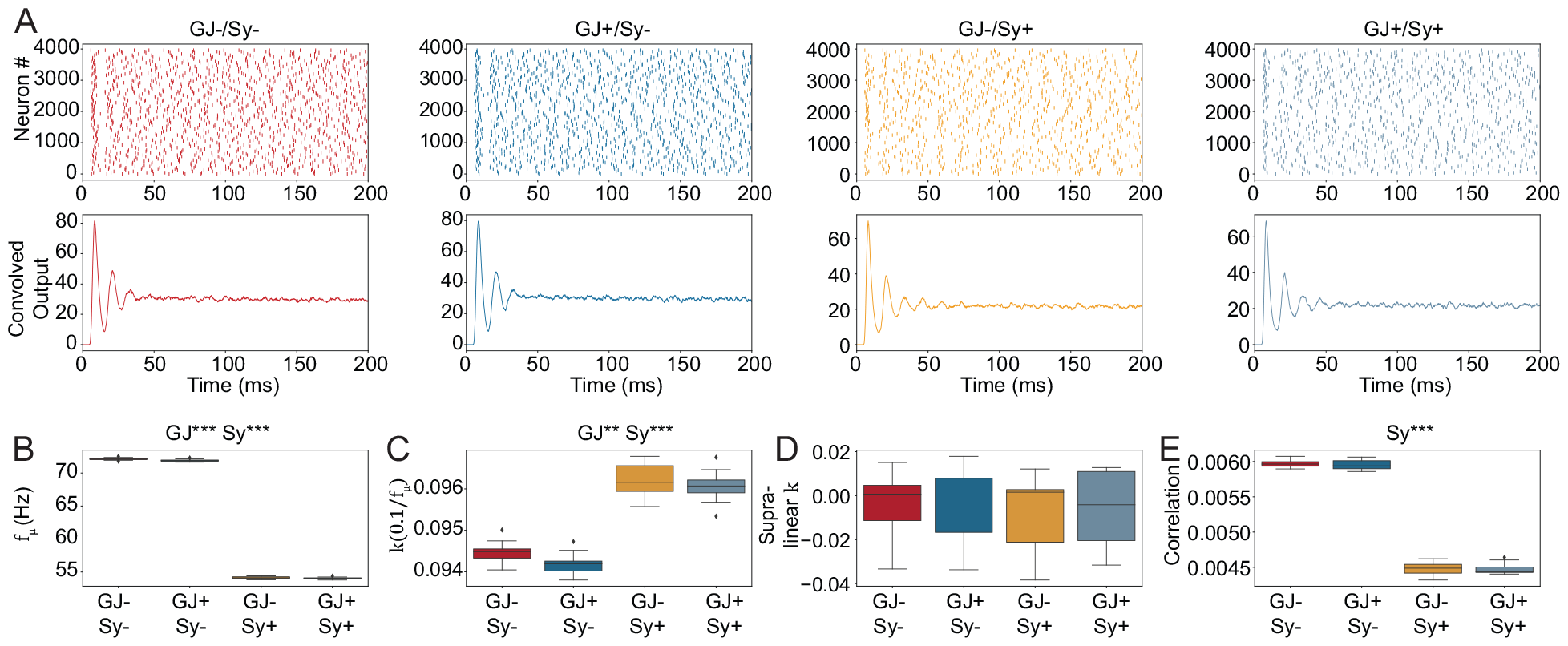
Large PV^+^ IN ring network. 4000 neurons were randomly distributed on a rectangular plane. **A** The spike raster plots show only every 40th neuron. Raster plot and convolved output show that the state is asynchronous. **B** Average frequency of all neurons in the network. Two-way ANOVA: Interaction: *F* = 1.893, *p* = 0.1743, Main effects: GJ, *F* = 14.0517, *p <* 0.001; Sy, *F* = 164105.4068, *p <* 0.001. **C-D** The synchrony measures. Two-way ANOVA for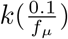: Interaction: *F* = 0.8884, *p* = 0.35, Main effects: GJ, *F* = 8.0496, *p <* 0.01; Sy, *F* = 540.5753, *p <* 0.001. Two-way ANOVA for Supralinear had no significant effects. Two-way ANOVA for Correlation: Interaction: *F* = 0.191, *p* = 0.6638, Main effects: GJ, *F* = 0.2828, *p* = 0.597; Sy, *F* = 6841.9982, *p <* 0.001. GJ, gap junction; Sy, synapse.

**Supp. Fig. 3:**
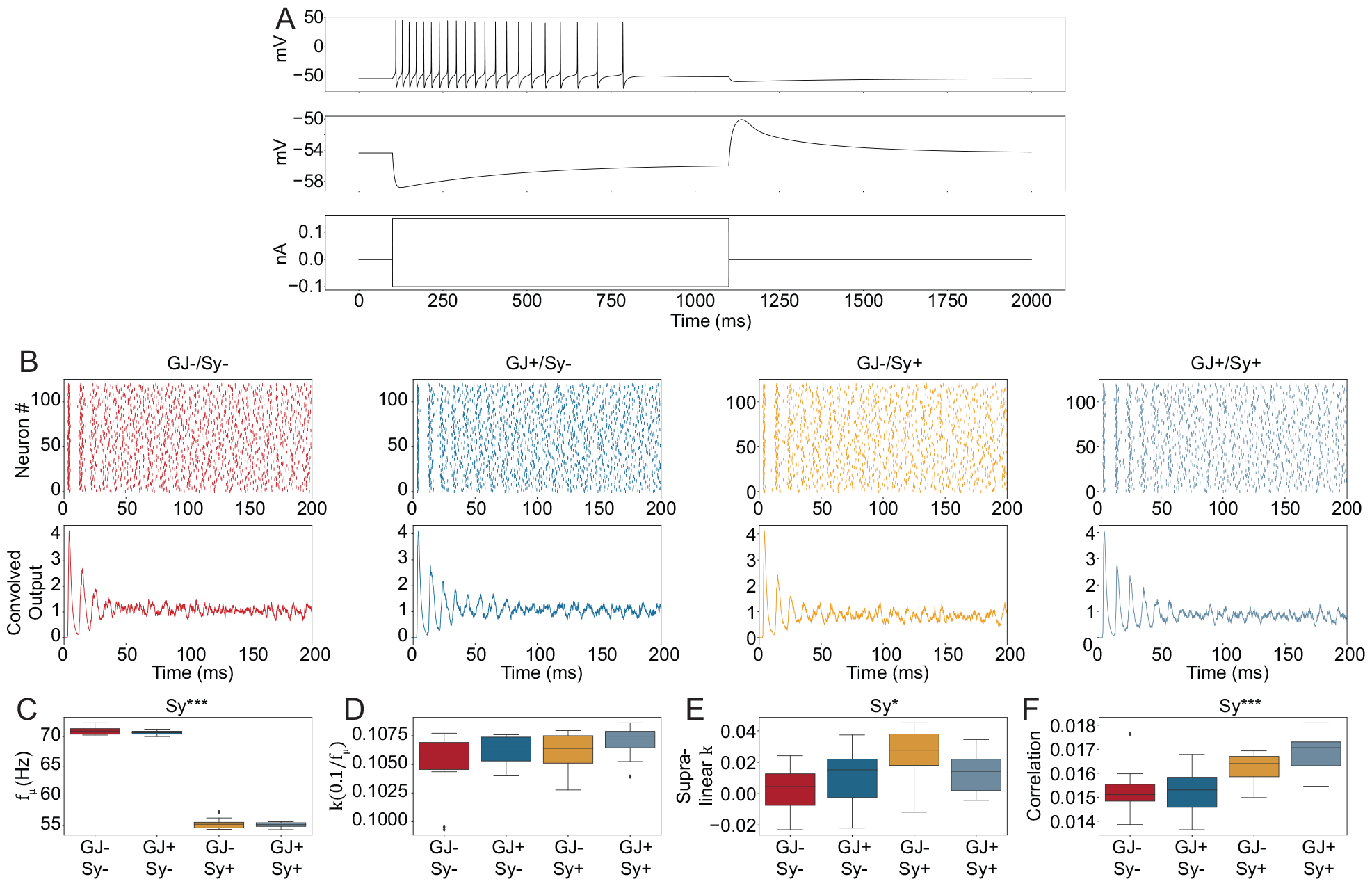
Ring network simulations with PV^+^ INs containing *I*_*h*_ current. **A** Voltage response of the modified PV^+^ IN with *I*_*h*_ added. Top: voltage response to depolarizing current step. Middle: response to hyperpolarizing current step with the slow depolarization characteristic for *I*_*h*_. Bottom: The current injections that induce the voltage responses above. **B** Spike raster plot and convolved output show an asynchronous state. **C** Average frequency of all neurons in the network. Two-way ANOVA: Interaction: *F* = 0.0926, *p* = 0.7626, Main effects: GJ, *F* = 1.8648, *p* = 0.1805; Sy, *F* = 5884.6182, *p <* 0.001. **D-F** The synchrony measures. Two-way ANOVA for 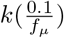 had no significant results. Two-way ANOVA for Supralinear k: Interaction: *F* = 4.0058, *p* = 0.0529, Main effects: GJ, *F* = 0.1206, *p* = 0.7304; Sy, *F* = 6.8143, *p <* 0.05. Two-way ANOVA for Correlation: Interaction: *F* = 1.2169, *p* = 0.2773, Main effects: GJ, *F* = 1.0703, *p* = 0.3078; Sy, *F* = 21.5984, *p <* 0.001.

**Supp. Fig. 4:**
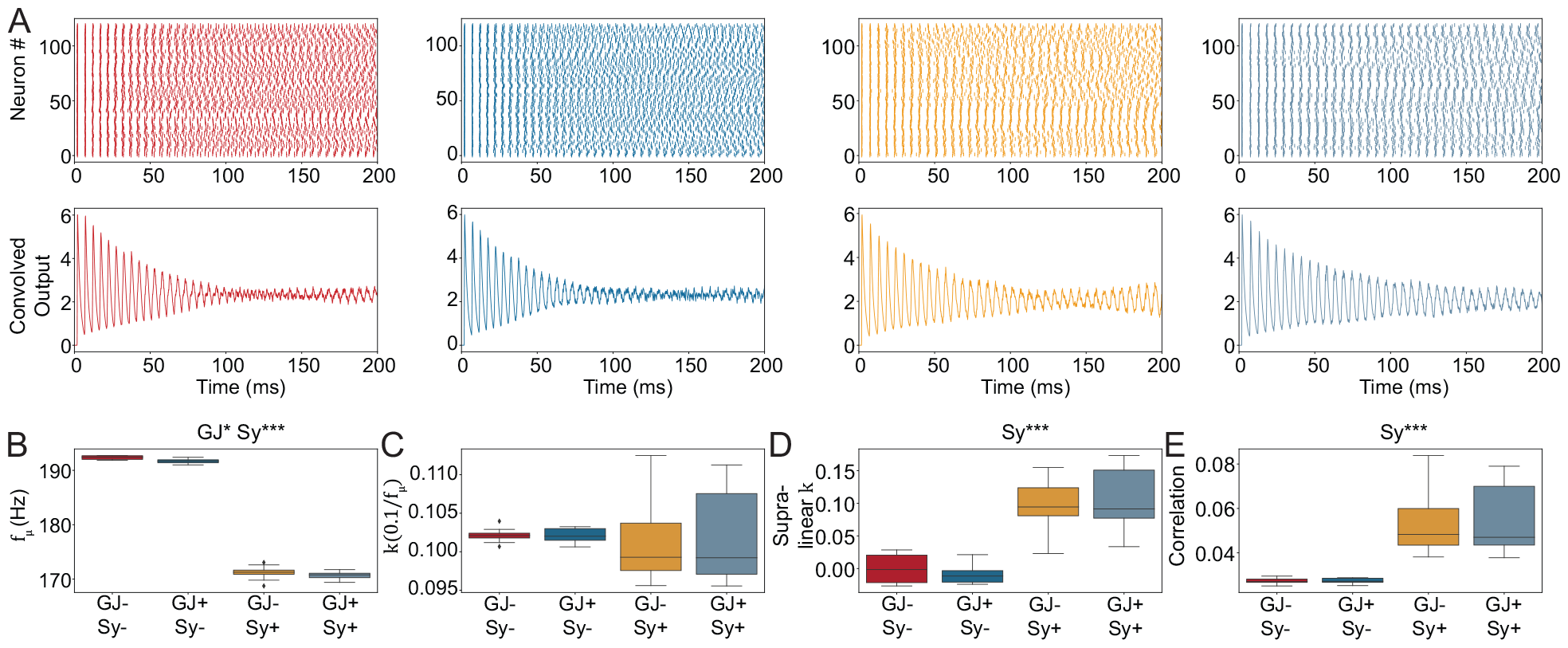
Ring network simulation with strong input current resulting in high average frequency. **A** Spike raster plot and convolved output show that some synchronous oscillations are larger when chemical synapses are added. However, this synchronous state is different from the one in **Figure 1**, where cells become silent during a cycle and furthermore does not reach the magnitude. **B** Average frequency of all neurons in the network. Two-way ANOVA: Interaction: *F* = 0.1127, *p* = 0.7391, Main effects: GJ, *F* = 5.3695, *p <* 0.05; Sy, *F* = 7474.6760, *p <* 0.001. **C-E** The synchrony measures. Two-way ANOVA for 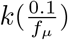 had no significant results. Two-way ANOVA for Supralinear k: Interaction: *F* = 0.9549, *p* = 0.3350, Main effects: GJ, *F* = 0.0117, *p* = 0.9145; Sy, *F* = 97.124, *p <* 0.001. Two-way ANOVA for Correlation: Interaction: *F* = 0.0059, *p* = 0.9393, Main effects: GJ, *F* = 0.0137, *p* = 0.9076; Sy, *F* = 60.3379, *p <* 0.001

**Supp. Fig. 5:**
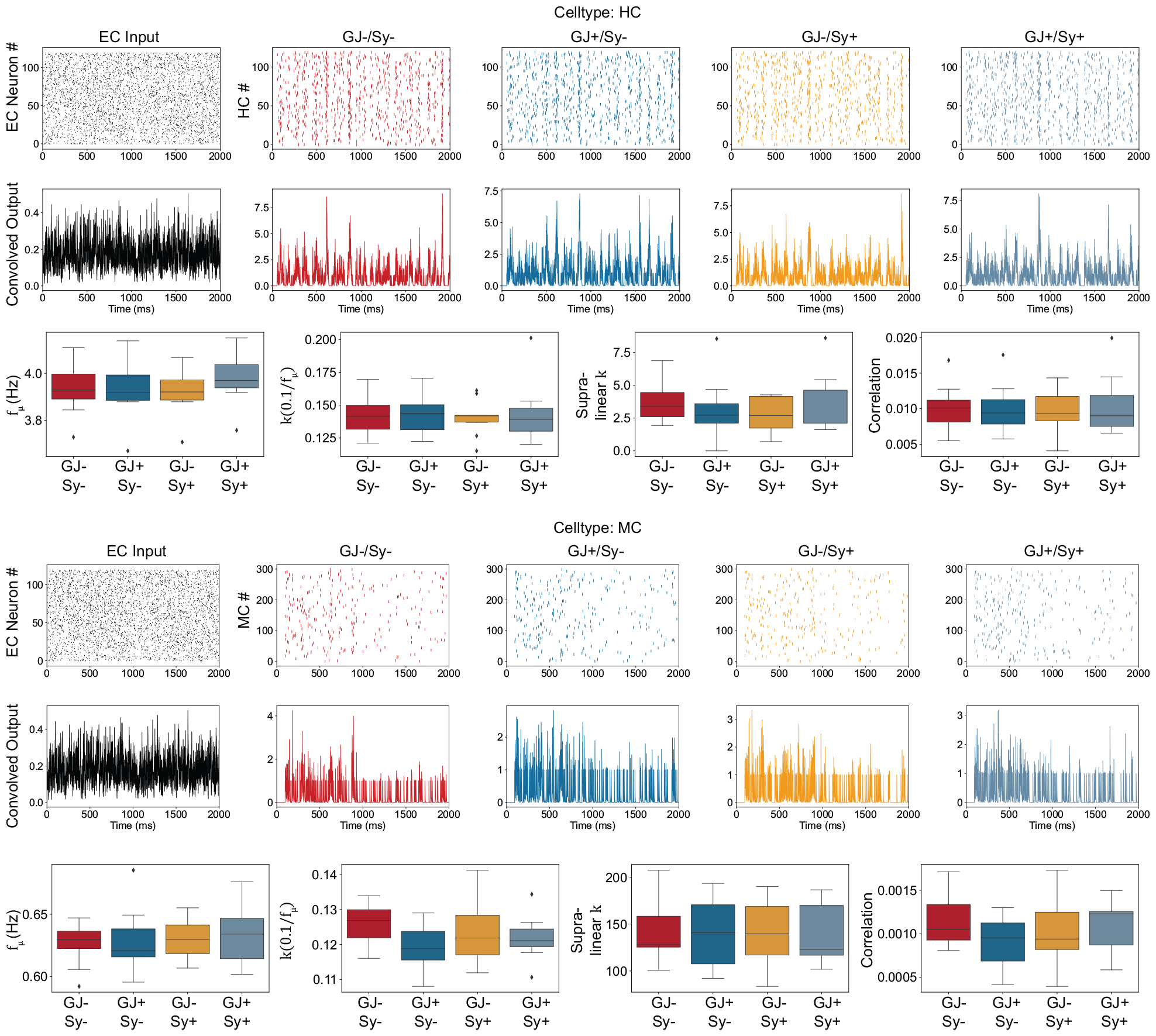
Results for the HC (top) and MC (bottom), which were simulated for **Figure 4** but not shown. Two-way ANOVA was done for all of the box plots but showed no significant result for any of them.

**Supp. Fig. 6:**
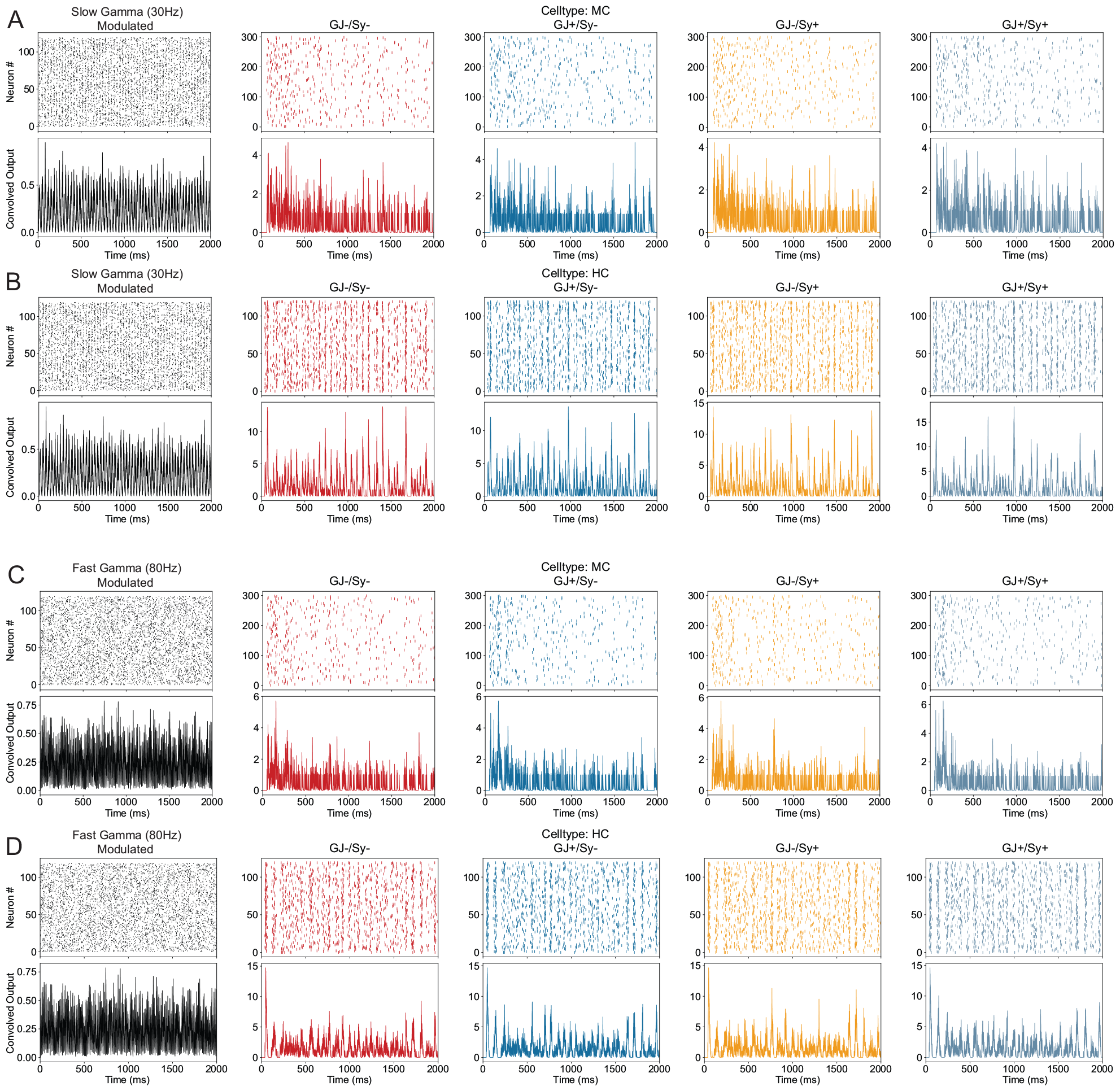
Cell types that were simulated in **Figure 5** but not shown. Boxplots were omitted because none of them was significant with the sole exception of a significant increase of average HC frequency in the fast gamma condition when synaptic connectivity is added in the PV^+^ ring network: *F* = 5.322589, *p <* 0.05.

## Notes

### Competing Interest Statement

The authors have declared no competing interest.

